# Likelihood of social-ecological genetic model

**DOI:** 10.1101/2025.03.06.641298

**Authors:** Stéphane Dupas, Camilo Patarroyo

## Abstract

The genetic structure of populations depends on two parallel processes - genetic and social-ecological - providing mutual information. Models that describe species’ responses within social-ecological systems are increasingly important in the context of modern environmental crises. Advances in genetic data collection now provide access to vast numbers of markers, enabling a more comprehensive understanding of species dynamics. However, current statistical inference methods do not fully integrate these processes into a single cohesive model, and rely on summary statistics derived from separate inferences and simulations.

In this work, I propose a probabilistic framework based on a coalescent model to compute the likelihood of a demogenetic model represented as connected graphs. The population graph, linking population nodes, is characterized by a backward gene adjacency matrix, which describes the probability of gene origins and is influenced by niche and dispersal functions. The genetic graph, linking allele nodes, captures mutation probabilities. A third graph represents the coalescence process of genes within the demographic and genetic graphs.

Coalescence is simulated within the demographic model, and its probability is estimated based on the genetic model, or vice versa. Likelihood estimation is performed using a Monte Carlo algorithm. This framework allow likelihood-based sampling and Bayesian inference, offering a robust approach for environmental niche and meta-population modeling.

I also discuss the practical applications of this model. By combining environmental niche functions with a coalescent framework, this approach enables probabilistic reconstructions of past species distributions based on present and historical occurrence data. It can incorporate various data sources, including historical records, absence data, and genetic information.

By integrating niche and dispersal processes into a unified model, this framework provides a powerful tool for improving species distribution forecasting and deepening our understanding of species’ responses to environmental change as interconnected systems.

**Author summary:** Inferring biodiversity responses to social-ecosystem history using population genetic and environmental history data is crucial in the context of modern environmental crises. However, existing models do not fully integrate key processes such as niche modeling, coalescent modeling, and genetic modeling, often relying on approximate Bayesian computation for inference. In this work, I propose a method to estimate the likelihood of a fully integrated demo-genetic model that links population genetic data with socio-ecosystem history.

The proposed method is based on a hidden coalescent process simulated within a social-ecosystem graph, along with a sampling process represented in a genetic mutation graph. In the social-ecosystem graph, nodes represent populations, and their effective sizes are estimated using niche models that account for environmental variables. The edges of the graph represent effective migration, which is estimated through connectivity models. The resulting likelihood function can be used for Bayesian or maximum likelihood inference of genetic changes within socio-ecosystems.

This approach offers a flexible framework for inferring demo-genetic models across various fields, including phylogeography, agrobiodiversity evolution, ecology, and epidemiology. It can be applied for forecasting or hypothesis testing, providing a comprehensive tool for understanding genetic dynamics in relation to socio-ecosystem processes.

## Introduction

In the context of the modern ecological crisis, understanding species’ responses to socio-ecosystem changes has become one of the most fundamental scientific challenges in ecology. Ecological niche theory is useful for predicting species’ responses; however, realized niches also depend on species interactions and historical dispersal events. Genes remain the fundamental unit of information transmission, making them particularly relevant for studying species’ responses in social-ecological systems [1]. The increasing availability of molecular data provides valuable complementary insights into realized niches.

One approach to integrating genetic information focuses on ‘omics, examining emergent properties from individuals to populations and ecosystems [2]. Other approaches emphasize population-level processes, both adaptive and neutral. Population genetic structure is shaped by adaptive (selection) and neutral forces (drift, mutation, migration). Many genetic and genomic studies now focus on documenting adaptive processes, particularly those related to global change and niche differentiation between populations [3]. A promising research direction is to consider neutral forces, including dispersal processes and historical events [4]. The presence of a gene at a given site is determined both by its environmental niche and its ability to reach that location through spatial and socio-ecological networks.

A significant advance in recent decades has been bridging the gap between population genetics and ecological niches. Niche models provide insights into past abundance and migration patterns. Drift is influenced by reduced population size and can be linked to environmental variables through niche functions. Existing models generally separate inference into two steps: first, estimating species distribution parameters from occurrence data, and second, inferring genetic evolution parameters from genetic data [5]. The term Integrated Distribution Demographic and Coalescence Model (IDDC) [6] was introduced to describe these efforts. Several simulation tools now allow genetic parameter inference within discrete or continuous species distribution landscapes [6–10]. Most of these models use niche modeling to simulate species distribution, construct evolutionary force landscapes, and perform coalescence simulations using Approximate Bayesian Computation (ABC). Many spatially explicit tools are available [9], with SPLATCHE [7, 11, 12] being one of the most advanced. This software enables the simulation of genetic data based on historical demographic maps and introduction scenarios. However, these IDDC approaches still present some limitations.

The first major limitation is the lack of full integration of niche, demographic, and genetic processes into a single model. In existing IDDC models, demographic parameter maps are not inferred but rather approximated using species distribution maps. There is no inference algorithm that simultaneously integrates demographic parameters, environmental relationships, and coalescent processes. Separating demographic and genetic inference is problematic because genetic drift does not depend solely on absolute population size but also on effective population size and migration rates [4]. Additionally, niche models used as proxies for historical reconstruction are typically based on occurrence or presence-absence data, which themselves depend on historical contingencies such as migration and extinction. Thus, there is no justification for treating niche and historical processes separately in demogenetic inference. Integrating niche and genealogical modeling would address these issues. QuetzalCoalT [13] is currently the only available tool capable of performing integrated inference of genetic and niche-based processes from environmental history and genetic structure. Recognizing that demographic and genetic processes should be fully integrated, we propose the term environmental demogenetics, as the ultimate inference model should incorporate environmental, demographic, and genetic data. Furthermore, most IDDC and environmental demogenetic models fail to integrate social-ecological networks, which are increasingly crucial in the Anthropocene era (e.g., agricultural practices, ethnic groups, trade routes).

The second major challenge of IDDC approaches is performing inference in the context of complex historical and high-dimensional processes. The relationship between environmental and genetic processes is typically a high-dimensional, unknown coalescent process. Consequently, models rely on simulations, with inference performed using ABC on summary statistics. ABC inference compares observed data with data generated by random simulators, making it particularly useful for Bayesian inference when likelihoods are intractable or highly complex. This is a common situation in population biology, where ecological and genetic data are the result of multiple unobservable processes—including genealogy, ecological interactions, genetic evolution, and sampling—that are highly interdependent and contingent. In ABC, priors are sampled based on summary statistics rather than exact likelihood calculations. However, this approach has drawbacks: information may be lost when transforming data into summary statistics, and there is no universal method for selecting appropriate summary statistics. Each inference problem requires a custom selection, and it is impossible to definitively prove that the chosen summary statistics capture all relevant aspects of the modeled process [14]. These limitations do not exist in likelihood-based inference.

To address these challenges, we propose a novel approach that:

1. Integrates social-ecological history, demographic processes, and genetic processes within a single inference model.
2. Directly incorporates multiple data sources, including genetic data, ecological occurrence data, and environmental history.
3. Accounts for both spatial and non-spatial networks, including social-ecological networks.
4. Performs exact likelihood-based probabilistic modeling, rather than relying on approximations.

Our metamodel consists of three interconnected graphs:

- A genetic graph, where nodes represent alleles and edges represent mutations.
- A population graph, where nodes represent populations and edges represent migration events.
- A coalescent graph, where nodes represent individuals and edges represent lineages.

The coalescent process serves as a fundamental link between genetic and demographic sampling processes. We propose two approaches for likelihood sampling within this metamodel using genetic status and demographic location data. First, coalescence can be simulated based on sample locations within the population graph, and the probability of the coalescent, the genetic data and graph model can be calculated. Alternatively, genealogy can be simulated within the genetic graph, and the probability of observed demographic locations can be calculated for the coalescent. This framework enables both likelihood-based and Bayesian inference. By integrating potential niche parameters (related to scenopoietic variables) with historical processes shaping the realized niche, this approach should enhance our understanding of species distributions and improve future projections (Figure 1).

**Figure 1.**
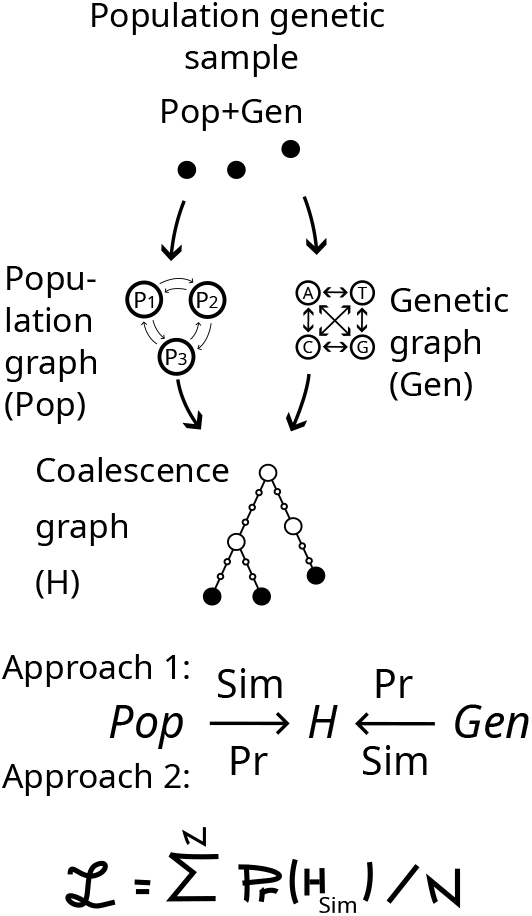
Social ecological demogenetic likelihood computation. Two approaches are considered. Approach 1: simulate coalescent from population information and calculate its probability from genetic information. Or Approach 2: simulate from genetic and calculate probability from populations. *Sim*: simulation of coalescent knowing one type of data and model. *Pr*: probability of coalescent knowing the other type of data and model.

### Demo-genetic Model

I present below the variables and parameters of the different graph models conforming the demogenetic model. The **population graph** represents populations, the **genetic graph** represents alleles and the **coalescence history graph** represent sample coalescing lineages (Table 2, figures 1,2,3).

**Table 1.** Definition of variables used in the model.

| Category | Symbol | Definition |
| --- | --- | --- |
| **Demographic Graph ($G_P$)** |  |  |
| | $V_P$ | Population (node) vector |
| | $V_P[i]$ | Population $i$ |
| | $t$ | Generation time |
| | $N_e[i, t]$ | Effective population size of population $i$ at generation time $t$ |
| | $X_e[i, t]$ | Vector of environmental variables for population $i$ at generation time $t$ |
| | $\theta_P^N$ | Parameters of the effective population size model |
| | $E_P$ | Backward population transition (edge) |
| | $M_P$ | Backward population transition matrix: probability that the father of an individual in population $i$ was from population $j$ |
| | $M_P[i, j]$ | Transition probability that an individual in population $i$ originated from population $j$ |
| | $m$ | Migration rate vector |
| | $m[i, j]$ | Probability that an individual from population $j$ migrates to population $i$ in the next generation |
| | $m(E_P[i, j], \theta_P^m)$ | Migration rate model |
| | $\theta_P^m$ | Migration rate model parameters |
| **Genetic Graph ($G_G$)** |  |  |
| | $V_G$ | Allele vector |
| | $V_G[i]$ | Allele $i$ |
| | $E_G$ | Backward allele transition data (edge) |
| | $M_G$ | Backward allele transition matrix: probability that an individual in allele $i$ originated from allele $j$ |
| | $M_G[i, j]$ | Probability that an individual in allele $j$ mutates into allele $i$ in the next generation |
| **Coalescent Graph ($H_C$)** |  |  |
| | $V_C$ | Vector of nodes in the coalescence graph |
| | $S$ | Sample vector |
| | $S_P$ | Deme of collection for $S$ |
| | $S_G$ | Alleles of $S$ |
| | $IN$ | Internal node vector |
| | $E_C[u, v]$ | Lineage edge from node $u$ to its coalescence ancestor $v = Adj(u)$ |
| | $p(In[i] \mid \theta_P)$ | Probability of a coalescence event given demographic models and data |
| **Full Coalescent Graph ($H_I$)** |  |  |
| | $I$ | Vector of individuals in the full coalescence history |
| | $t(I)$ | Generation times of individuals $I$ in $H_I$ or $H_C$ |
| | $An(I, t)$ | Set of ancestors of individuals $I$ at time $t$ (duplicates removed) |
| | $P_P(I, t)$ | Matrix of probability distribution of $An(I, t)$ across demes in $G_P$ at time $t$ |
| | $P_P(I)$ | Matrix of demographic distribution of $I$ at $t(I)$ |
| | $P_G(I, t)$ | Matrix of probability distribution of $An(I, t)$ across alleles in $G_G$ at time $t$ |
| | $P_G(I)$ | Matrix of genetic distribution of $I$ at $t(I)$ |

**Table 2.** Signification of the variables.

|  |  |  |
| --- | --- | --- |
| **Demographic** | $G_P$ | **Demographic graph** |
| | $V_P$ | Population or node vector |
| | $V_P[i]$ | Population i |
| | $Ne$ | Effective population size vector |
| | $t$ | Generation time |
| | $Ne[i, t]$ | Effective population size of population i at generation time t |
| | $Xe[i, t]$ | Vector of environmental variables of population i at generation time t |
| | $\theta_P^N$ | Effective population size model parameters |
| | $E_P$ | Backward population transition or edge data variable |
| | $M_P$ | Edge Backward population transition matrix that individual in population i comes from population j |
| | $M_P[i, j]$ | Backward population transition probability that individual in population i comes from population j |
| | $m$ | Migration rate edge vector |
| | $m[i, j]$ | Probability that an individual belonging to population j migrates to population i in the next generation |
| | $m(E_P[i, j], \theta_P^m)$ | Migration rate model |
| | $\theta_P^m$ | Migration rate model parameter |
| **Genetic** | $G_G$ | **Genetic graph** |
| | $V_G$ | Allele vector |
| | $V_G[i]$ | Allele i |
| | $E_G$ | Backward population transition or edge data variable |
| | $M_G$ | Edge Backward allele transition matrix that individual in allele i comes from allele j |
| | $M_G[i, j]$ | Probability that an individual belonging to allele j mutates to allele i in the next generation |
| **Coalescent** | $H_C$ | **Sample coalescence history graph** |
| | $V_C$ | Coalescence graph node vector |
| | $S$ | Sample vector |
| | $S_P$ | Deme of collection of $S$ |
| | $S_G$ | Alleles of $S$ |
| | $IN$ | Internal nodes vector |
| | $E_C[u, v]$ | lineage egce from a node $u$ to its coalescence ancestor $v = Adj(u)$ |
| | $p(In[i] \mid \theta_P)$ | the probability of a coalescence event knowing demographic models and data. |
| **Full coalescent** | $H_I$ | **Individual sample coalescence history graph** |
| | $I$ | Vector of individual nodes in full coalescence history |
| | $t(I)$ | generation times of individuals $I$ in the graph $H_I$ or $H_C$ |
| | $An(I, t)$ | The set of ancestors at time $t$ of the set of individuals $I$ (doublons being removed) |
| | $P_P(I, t)$ | Matrix of probability distribution of $An(I, t)$ in line among demes in $G_P$ in column, at ancestral time $t$. |
| | $P_P(I)$ | Matrix of demographic distribution of $I$ at $t(I)$. |
| | $P_G(I, t)$ | Matrix of probability distribution of $An(I, t)$ in line among alleles in $G_G$ in column, at ancestral time $t$. |
| | $P_G(I)$ | Matrix of genetic distribution of $I$ at $t(I)$. |

**Figure 2.**
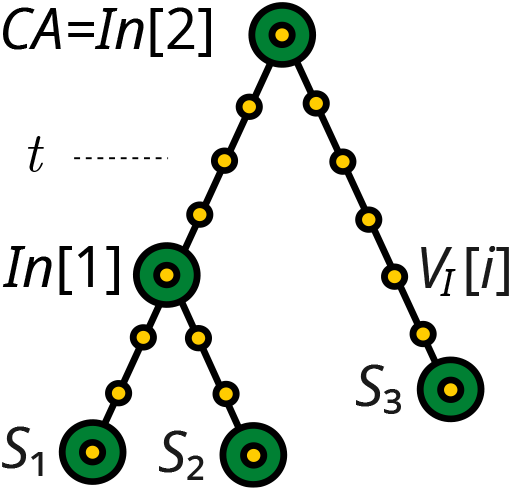
The coalescent graph *H*_*C*_ (green nodes) and the individuals coalescent graph *H*_*I*_ (yellow nodes). *S*: Sampled individuals. *In*: Internal coalescence nodes. *V*_*I*_ [*i*]: Nodes representing individuals within to the coalescent lineages. *H*_*C*_ = (*V*_*C*_, *E*_*C*_) consists of vertices *V*_*C*_ = *S* ∪ *In* and edges, *E*_*C*_, which define the lineages connecting them. The individuals along these edges belong to *H*_*I*_. *t*: generation time.

**Figure 3.**
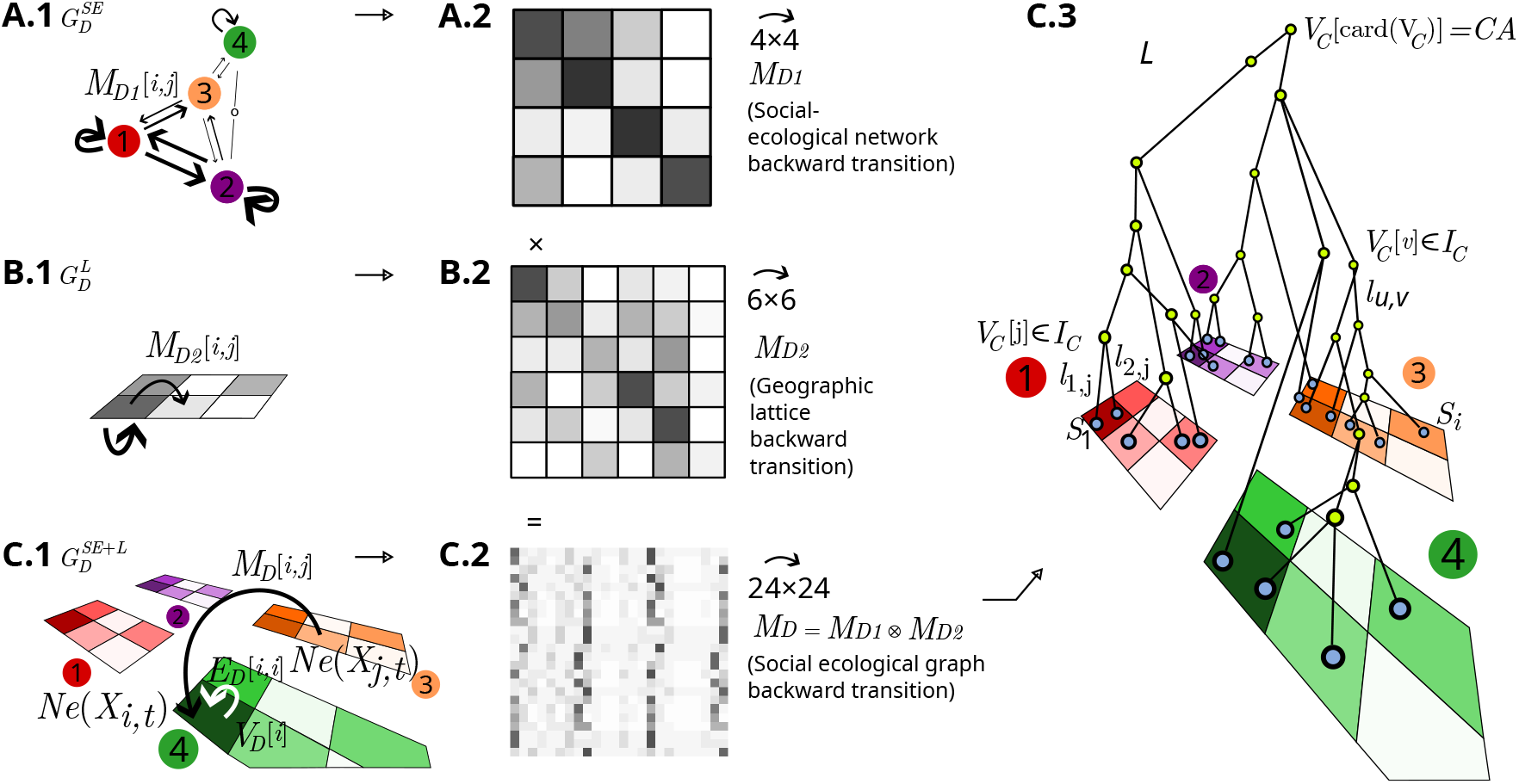
Examples of population graphs and a coalescence history graph simulated within a population graph. **(A.1)** Social-Ecological Network Demographic Graph 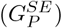 with 4 nodes, 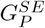.**(B.1)** Landscape Demographic Graph 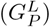 with 6 nodes, where nodes correspond to raster cells or vector map contours. **(B.2)** Backward population transition matrix among nodes in 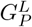.**(C.1)** Combined Social-Ecological and Landscape Demographic Graph 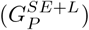,integrating 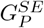 and 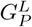. *V*_*P*_ [*i*]: Node (deme) *i. Xe*_*i*_: Environmental variables associated with node *V*_*P*_ [*i*]. *Ne*(*X*_*i*_): Effective population size of node *V*_*P*_ [*i*]. **(B.3)** Construction of the 24 × 24 backward gene population transition matrix *M*_*P*_, characterizing the edges 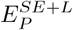 of 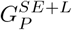 for coalescence simulations. **(C)** Movement and coalescence of a sample *S* = {*S*_1_, *S*_2_, …} within 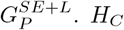 represents the coalescence history graph on 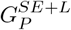,where *H*_*C*_ = (*V*_*C*_, *E*_*C*_) consists of vertices *V*_*C*_ = *S* ∪ *I*_*C*_ (with *I*_*C*_ as internal nodes of *V*_*C*_) and edges *E*_*C*_, representing the lineages connecting them.

**The population graph**, **noted** *G*_*P*_ = (*V*_*P*_, *E*_*P*_).

I first consider demographic history in a general random walk across a population graph model, representing populations as nodes (in spatial or ecological dimensions) and migration as edges (Figure 3). The population graph can represent a geographic landscape 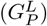,a socioecological network 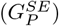,or a combination of both 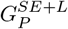 (Figure 3). The variables characterizing populations can also vary over time. Additionally, the migration function can depend on space and environment.(Figure 3).

- Each node *V*_*P*_ [*i*] represents a distinct population with an effective population size *Ne*[*i*].
- Each edge *E*_*P*_ [*i, j*] is characterized by a backward gene transition probability *M*_*P*_ [*i, j*], which represents the probability that a gene in node *V*_*P*_ [*i*] originated from node *V*_*P*_ [*j*] in the previous generation. This probability depends on the effective population sizes of both the receiving and originating demes *N*_*e*_, and the migration rate *m*[*i, j*], which is the proportion of individuals migrating from node *j* to node *i* in forward time.
- The effective population size *Ne* is defined by a niche model 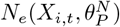,
- which relates *Ne* to environmental variables
- *Xe*_*i,t*_ and model parameters 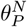.
- The migration rate m[i,j] is defined by a migration rate model *m*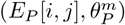,which relates m to a distance measure between nodes *i* and *j*, incorporating migration-friction parameters 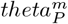.
- The number of demes (the order of the graph) is denoted as |*D*|, while the number of migration connections (the degree of the graph) is denoted as |*E*_*P*_ |. The matrix *M*_*P*_ represents the backward transition probabilities between populations.

#### The genetic connected graph, denoted as *G*_*G*_ = (*S*_*G*_, *E*_*G*_)

We now consider the coalescence in the genetic graph, where nodes represent alleles and edges represent mutations.

- Each node *V*_*G*_[*i*] represents an allele state.
- Each edge *E*_*G*_[*i, j*] represents a mutation event that is characterized by the probability *M*_*G*_[*i, j*], that a gene in node *V*_*G*_[*i*] have mutated from node *V*_*G*_[*j*] in the previous generation. The matrix *M*_*G*_ represents mutation rate between alleles.
- The number of alleles (the order of the graph) is noted |*V*_*G*_| while the number of mutation connections (the degree of the graph) is denoted |*E*_*G*_|.

**The coalescence history graph, denoted** *H*_*C*_(*S*):

*H*_*C*_(*S*) is an directed graph that traces the lineage of a genetic sample *S* back to their common ancestor, which edges oriented opposite to those in a traditional genealogical graph (Figure 2).

- Each node *V*_*C*_[*i*] represents either a sample *S* or a coalescence point.
- Each edge *E*_*C*_[*u, v*] represents a lineage connecting a node to its next coalescence point.
- The coalescence history graph is defined as HC=(VC,EC) *H*_*C*_ = (*V*_*C*_, *E*_*C*_), where *E*_*C*_ = {(*u, v*) ∈ *V*_*C*_, where v is parent of u}).
- Sampled individuals *S*_*i*_ = *V*_*C*_[*i*] with *i* = {1..*n*} are defined as nodes with zero in-degree. These individuals are linked to their ancestors through coalescence events, which correspond to internal nodes *In*. Internal nodes are characterized by having more than one in-degree and exactly one outdegree, except for the most recent common ancestor (MRCA), which has more than one in-degree and zero out-degree.

**The individual colaescence graph, denoted** *H*_*I*_ (*S*) (Figure 2). The individual colaescence graph *H*_*I*_ (*S*) is an extended graph that includes the coalescence history graph and all individuals along the edges of the coalescence history graph that constitute its lineages. It is defined as *H*_*I*_ = (*I, E*_*I*_). Such that *V*_*C*_ ⊂ *I*, and *E*_*I*_ = {(*u, v*) | *v* is the parent of *u* in the previous generation}

- Each node represents an individual kin the genealogy of the sampled genes. It is either a node from *H*_*C*_(*S*), or an individual positionned along the vertices *V*_*C*_(*S*) between two connected nodes of *H*_*C*_.
- *H*_*I*_ (*S*), as *H*_*C*_(*S*) is an acyclic graph oriented from the sampled individuals to the common ancestor. Each individual *I*[*i*] ∈ *I* is has a unique parent formally expressed as card(*Adj*(*I*[*i*]) = 1. The descendants of *I*[*i*] are obtained by *Adj*^−1^(*I*[*i*]).
- Each individual of *H*_*C*_ is fully defined within the coalescent tree by one of its descendant, including sampled descendant *S*_*i*_ and the generation time (Figure 2).

The probability of a coalescence history graph *H*_*C*_ depends on both the genetic graph and the population graph. To estimate the likelihood of the demogenetic model given environmental, demographic, and genetic data, we will simulate coalescence history graphs within the population graph and compute their probabilities under the genetic graph.

All the variables and terms used in this paper are described in the table 2.

### Simulation of genealogy knowing demographic model

The first method of likelihood calculation consists of simulating a genealogy of the sample in a population graph, and then evaluating its probability in the genetic graph. The species observation data used in the model may include genetic information on the individuals reported, but this is not always required.

To analyze the movements and coalescence of gene lineages within the population graph, we first compute the **population transition matrix**, *M*_*P*_. This matrix is a Markov transition matrix that estimates the evolution of occurrence probabilities among demes. Each cell [*i, j*] represents the backward in-time probability, that a gene in deme (or node) *i* originates from deme (or node) *j*.

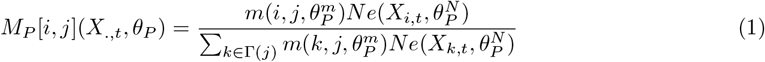

Where Γ(*i*) represents the set of all demes directly connected to 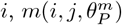 is the probability function of migration from deme *i* to deme *j* according to the migration model 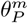,and 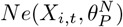 is the effective population size given by a niche model that depends on environmental variables and parameters. Note that 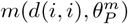 corresponds to the probability that genes remain in the same deme.

Additionally, the matrix satisfies the condition ∑_*j*_ *M*_*P*_ [*i, j*] = 1, ensuring that transition probabilities sum to one.

We will consider coalescence events sequentially, from the most recent sampling date back to the MRCA, proceeding by pairs of lineages. To represent coalescence nodes, we observe that the function *An*(*S*[*i*], *t*) is an injection from *S* × *t*(*I*) to *I*: each individual can be uniquely identified in *G*_*I*_ by one of its descendant, in particular in the sample, and its generation time.

Note that if coalescence event occur at time *t* between two lineages *An*(*I*[*i*], .) and *An*(*I*[*j*], .), then for any earlier time *τ* such that *τ* ≤ *t*, it follows that: ⇒ *An*(*I*[*i*], *τ*) = *An*(*I*[*j*], *τ*).

If no coalescence occur, the evolution of the, matrix *P*_*P*_ (*S, t*) from time *t* to *t* − 1 is given by :

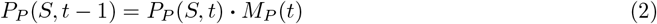

which describes the transition between generations

Similarly, if no coalescence occurs, the general term at time *t* is :

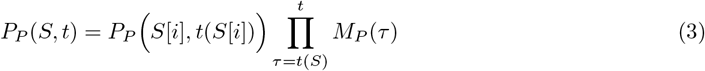

Consider two ancestors *An*(*S*[*i*], *t*) and *An*(*S*[*j*], *t*) that have not yet coalesced at generation *t*. The probability that they coesist in the same deme in the previous generation *t* − 1 is given by the scalar product of their population distribution probabilities at generation *t*:

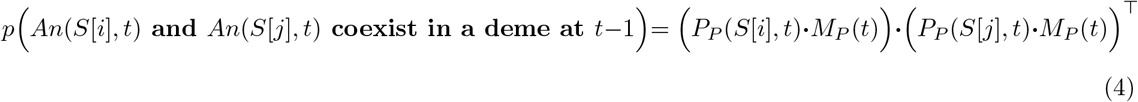

The probability that they coalesce in the previous generation (*t* − 1) is obtained by dividing the vector of coexistence probabilities across demes by twice the effective population size predicted by the niche model in each corresponding deme *N*_*e*_(*X*(*t* − 1)):

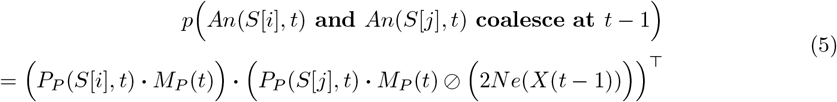

Where ⊘ is the element-wise division operator.

If two lineages coalesce, the probability distribution of their common ancestor given the demographic model, is given by performing an element-wise multiplication of the distribution probabilities of the two lineages, followed by an element-wise division by the population size. The result is then normalized by the probability of coalescence.

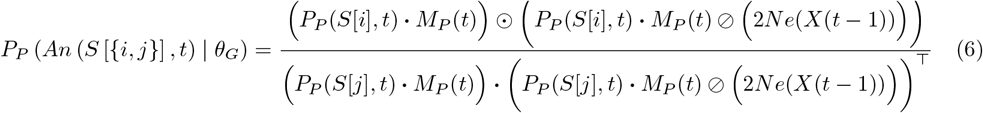

Where:

**bla** ⊙ represents element-wise

**bla** ⊘ represents element-wise division.

Using these equations of genetic distribution and coalescence probability from one generation to the parent, we can simulate their genealogy using the following algorithm.

#### Algorithm 1

Demographic simulation of a coalescent

1. **Initialize generation time:** Define t = max{t | card(An(S, t)) < 2}, i.e., the most recent time when at least two lineages exist.
2. **Compute the genetic distribution:** Calculate the genetic distribution PP (S, t) using equation (3).
3. **Simulate coalescence events in pairs:**

(a) Randomly select a pair of lineages in the set An(S, t).
(b) Simulate their coalescence at t – 1 with the probability defined in equation (5).
4. **Update lineages if coalescence occurs:**

(a) If the selected pair coalesces, merge their lineages.
(b) Update their demographic distribution using equation (6).
(c) Record the event in a genealogy object, including the probability distribution of each remaining lineage.
5. **Repeat for all remaining pairs:** Return to step 3 until all pairs in An(S, t) have been tested for coalescence.
6. **Move to the previous generation:** Decrease the time by setting t = t ߝ 1.
7. **Repeat until complete coalescence:** Return to step 2 until all lineages have merged into a single ancestor.

## Probability of a genealogy knowing genetic model

In the first approach, we consider a coalescent simulated under a known demographic model, as described in Algorithm 1.

Let us consider a coalescent graph *H*_*C*_ simulated under a given demographic model. We calculate the probability of each coalescence point under genetic model *p*(*In*[*k*] | *θ*_*G*_), characterized by *t*(*In*[*k*]) = *t*_*C*_ and its descendant nodes denoted as *In*[*k*]^*d*^ = *Adj*^−1^(*In*[*k*]). The number of descendant lineages coalescing at *In*[*k*] is given by *In*[*k*]^*Nd*^ = card(*In*[*k*]^*d*^) = card(*Adj*^−1^(*In*[*k*])). This number can be 2 in the case of a dichotomy or greater in the case of a polytomy. These coalescence events occur at times *t*(*In*[*k*]^*d*^).

If the lineages have not coalesced at any more recent time, the probability that they co-occur in the same genetic state (i.e., in the same population) at *t* | *t* ≤ min *In*[*k*]^*D*^, given the genetic model *θ*_*G*_, is given by the scalar product of the probability distributions of each individual in *In*[*k*]^*d*^ at time *t* (obtained from equation (3) applied to individuals in *In*[*k*]^*d*^).

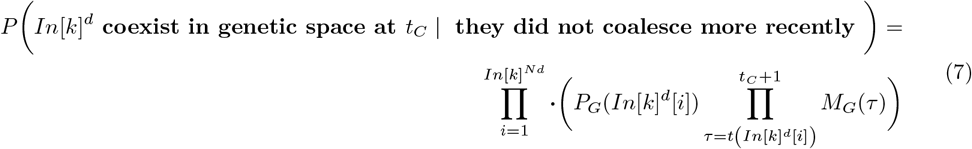

Where “∏ ·” represents the scalar product of the vectors.

The probability of coalescence is proportional to this probability (as it depends on demographic data and model, and is independent on genetic data and model). If the individuals have not coalesced at a more recent generation, the probability that two individuals coalesce at generation *t*_*C*_ = *t*(*In*[*k*]) is given by the scalar product of their probability distributions at generation *t*_*C*_, multiplied by the parameter 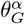 (since 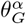 is independent of generation time):

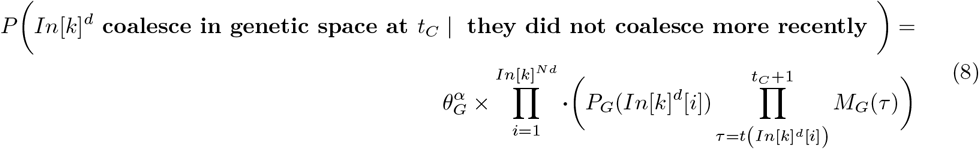

The parameter *θ*^*α*^ will be estimated as a coefficient related to the average effective population size *N*_*e*_.

The probability that they have not coalesced more recently is given by Equation **??**:

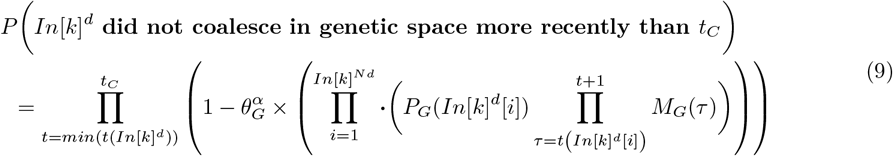

The final probablity of coalescence is :

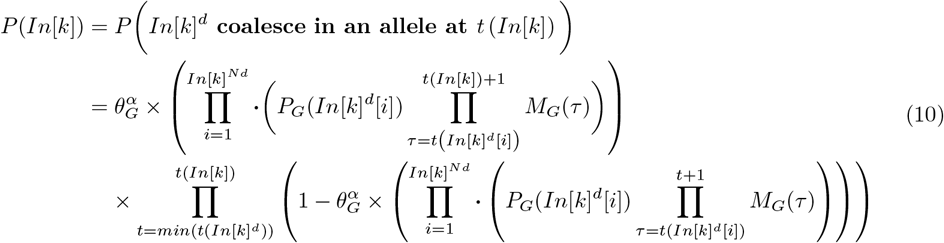

The allele probability distribution vector of the coalescent ancestor *In*[*k*] is given by:

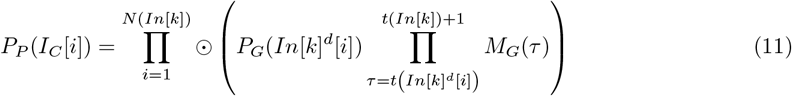

Where ∏ ⊙ is the element-wise vector multiplication.

We can calculate the probability of a data sample (*P*_*G*_(*S*) or *S*_*G*_) and genealogy *H*_*C*_, given the genetic model *θ*_*G*_, by multiplying the probabilities from Equation (**??**) over coalescence events and updating the probability distribution of nodes across the genetic space using the corresponding equation:

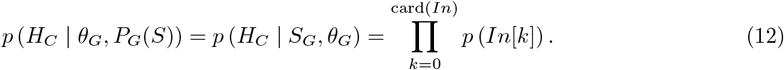

### Algorithm 2

Genetic Probability of a Demographically Simulated Coalescent

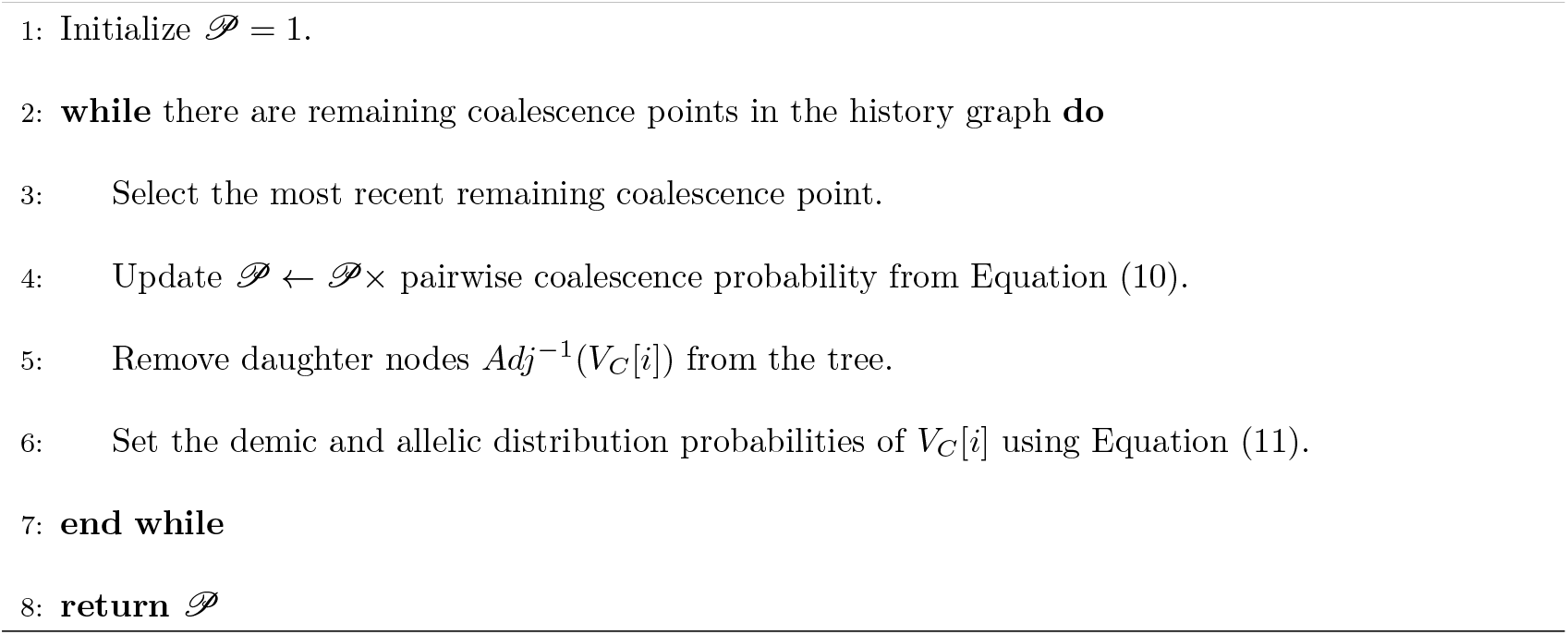

Note that this algorithm 2 can be combined with the demographic coalescence simulation algorithm (Algorithm 1) so that the coalescence simulation yields directly the genetic probability of the demographic coalescent.

**Algorithm 3: Joint simulation of coalescent on population graph and calculation of the probablity on genetic graph**

1. Set generation time *t* = {max *t* | card(*An*(*S, t*)) ≥ 2}. Set *P*_*G*_ = 1
2. Calculate genetic distribution of *P*_*P*_ (*S, t*) (eq. 3).
3. Select randomly a pair of lineage in the set *An*(*S, t*) that remain. Simulate their coalescence or not at *t* − 1 with probability defined in eq. 5.
4. if coalescence occurred, merge the lineages, update demographic distribution using eq. 6. Record the event in a genealogy object including the probablity distribution location of each remaining lineage. Reset current *P*_*G*_ multiplying it by the pairwise coalescence probability of this coalescence event defined in (equation 10). Set the demic and allelic distribution probabilities of *V*_*C*_[*i*] acording to equation (11).
5. go to 3. until every pair of the set has been tested for coalescence.
6. Set *t* = *t* − 1.
7. go to 2. Until all lineages have coalesced

### Algorithm 3

Joint Simulation of Coalescent on Population Graph and Calculation of Probability on Genetic Graph

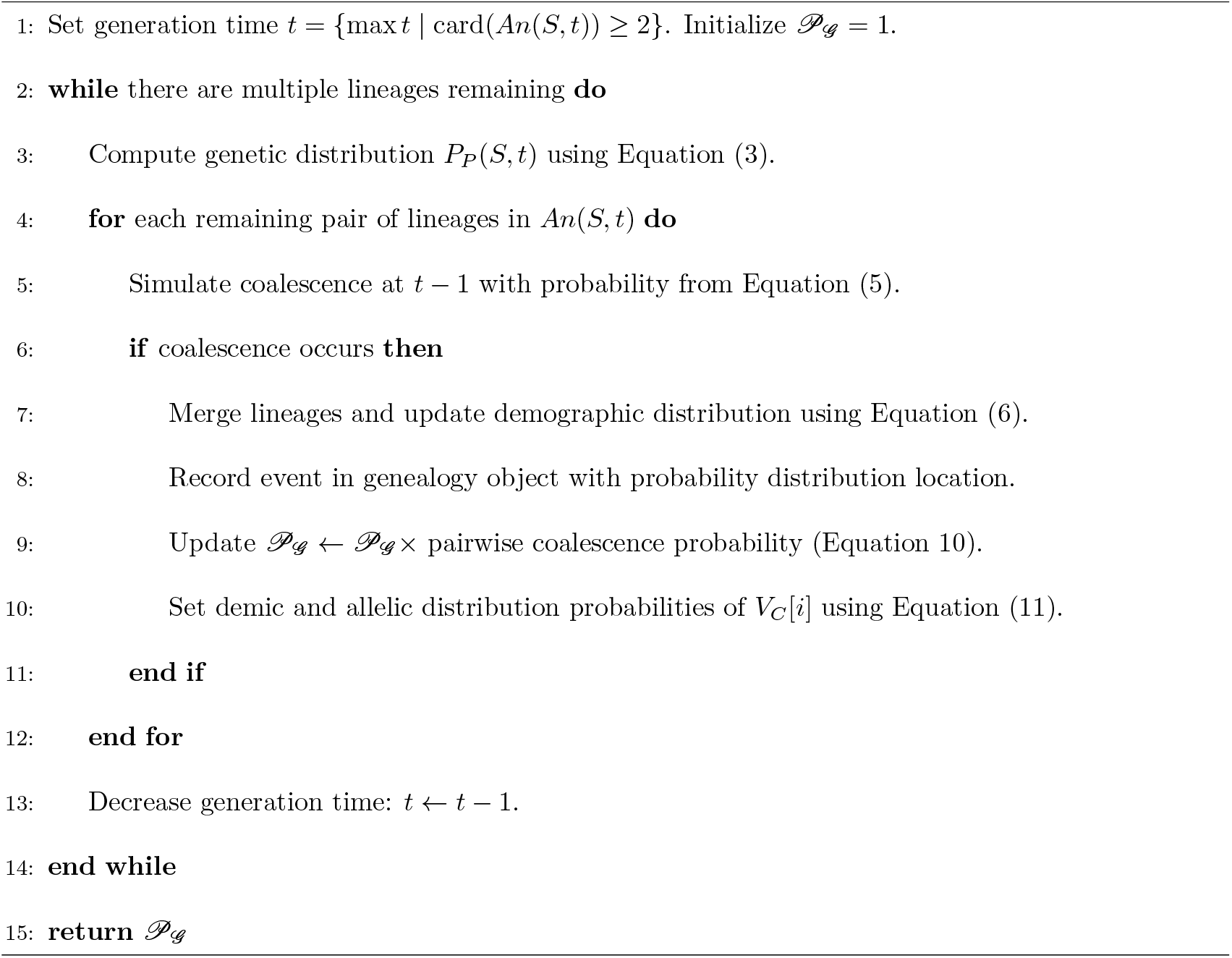

### Simulation of genealogy knowing genetic model

The second method of likelihood calculation consists of simulating a genealogy of the sample within the genetic graph and then evaluating its probability in the population graph. We again consider coalescence events from the most recent ancestor to the common ancestor, proceeding in pairs of lineages.

The same equations, namely Equation (2), Equation (3), and Equation (4), can be used to calculate the probability of transition from the genetic distribution at time *t* to time *t* − 1 (Equation 13), the general expression of the genetic distribution at any time *t* (Equation 14), and the probability that two genes, which have not coalesced more recently, share the same genetic state at ancestral time *t* (Equation 15), respectively.

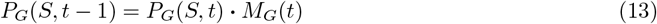

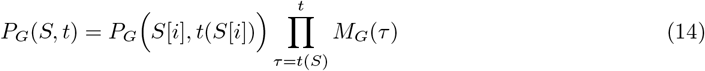

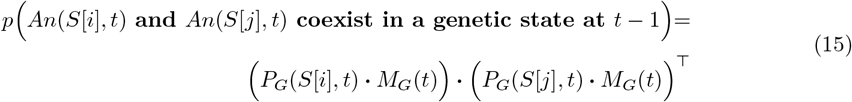

The probability that two lineages coalesce in the previous generation, *t* − 1, is obtained by multiplying the vector of coexistence probability in genetic space by a parameter 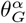,This parameter characterizes the probability of coalescence for individuals sharing alleles. It reflects the average effective population size, which is independent of the other *θ*_*G*_ parameters and *P*_*G*_(*S*) and, therefore, remains constant. However, this parameter is unknown and must be estimated.

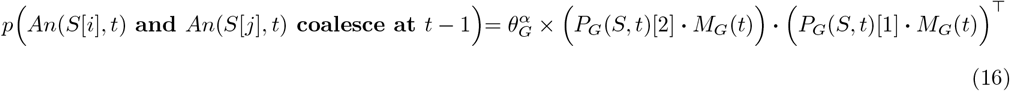

If they coalesce, the genetic distribution of the ancestor knowing genetic model model is given by elementwise multiplication of the distribution probability of the two lineages divided by the probability of coalescence (eq. 16). We can simplify by 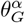 and obtain:

If coalescence occurs, the genetic distribution of the ancestor, given the genetic model, is obtained by performing an element-wise multiplication of the probability distributions of the two lineages and normalizing by the probability of coalescence (Equation 16). Simplifying by 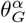,we obtain:

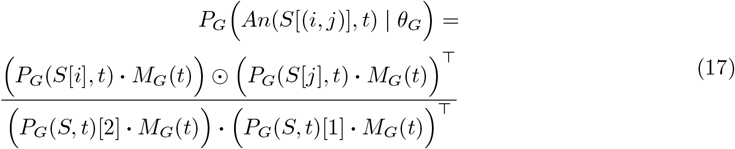

Using these equations for genetic distribution and coalescence probability from one generation to the parent, we can simulate their coalescence using the following algorithm.

#### Algorithm 4

Simulation of the genealogy given genetic graph and data

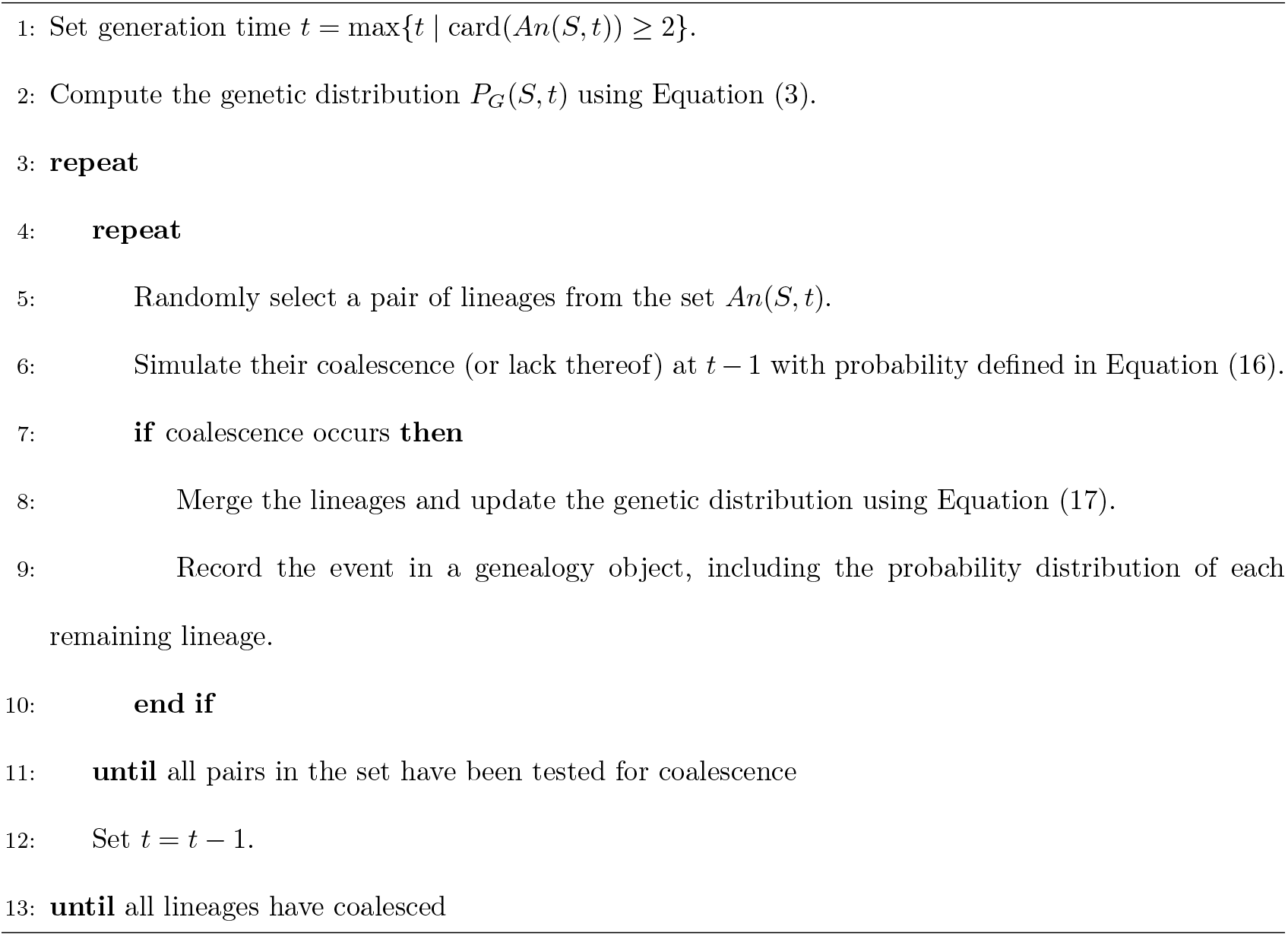

### Probability of a genealogy knowing demographic model

Let us consider a coalescent graph *H*_*C*_ simulated knowing the genetic model. We calculate the probability of the genealogy given the demographic model for each coalescence point *p*(*In*[*k*] | *θ*_*P*_), characterized by *t*(*In*[*k*]) = *t*_*C*_ and its descendants, denoted as *In*[*k*]^*d*^ = Adj^−1^(*In*[*k*]). The number of descendants coalescing in *In*[*k*] is given by card(*In*[*k*]^*d*^) = card(Adj^−1^(*In*[*k*])).

The number of descendants can be **2 in case of dichotomy**, or **more in case of polytomy**.

These events occur at time *t*(*In*[*k*]^*d*^).

If they do not coalesce more recently, the probability that they co-occur in the same genetic state (in the same population) at *t* (where *t* ≤ min (*In*[*k*]^*D*^)) given the demographic model *θ*_*P*_ is obtained by computing the scalar product of the probability distributions of each individual at time *t*, as given by Equation 3.

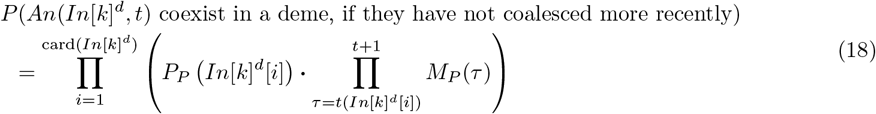

Let us consider a coalescent graph *H*_*C*_ simulated knowing genetic model. We calculate the probability knowing demographic model for each coalescence point *p*(*In*[*k*] | *θ*_*P*_) characterized by *t*(*In*[*k*]) = *t*_*C*_ and its descendants noted *In*[*k*]^*d*^ = *Adj*^−^1(*In*[*k*]). The number of descendants coalescing in *In*[*k*] is noted card(*In*[*k*]^*d*^) = card(*Adj*^−^1(*In*[*k*])). They can be 2 in case of dichotomy, or more in case of polytomy. They occur a times *t*(*In*[*k*]^*d*^)

If they do not coalesce more recently, the probability that they co-occur in the same genetic state (in the same population) at *t* ((*t* ≤ *min In*[*k*]^*D*^ )) knowing demographic model *θ*_*P*_ is given by the scalar product of the probability distributions of each individual at time *t* (given by eq. 3).

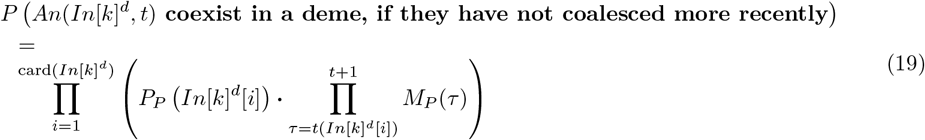

To obtain the probability that they coalesce at time *t*, we divide for each deme the probability of coexistence by the effective population size of each deme before computing the same scalar product:

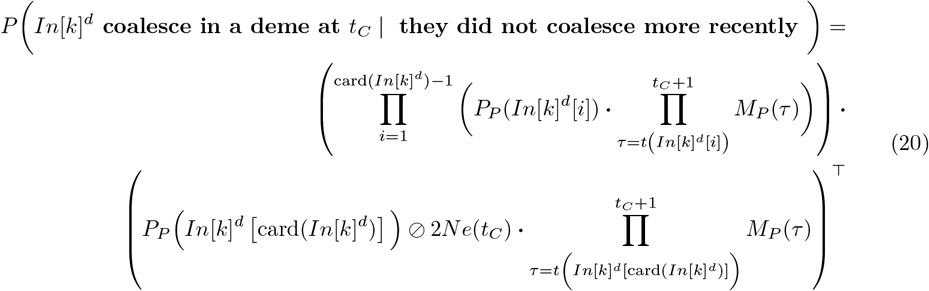

We now consider the probability that they **did not coalesce** more recently than time *t*_*C*_:

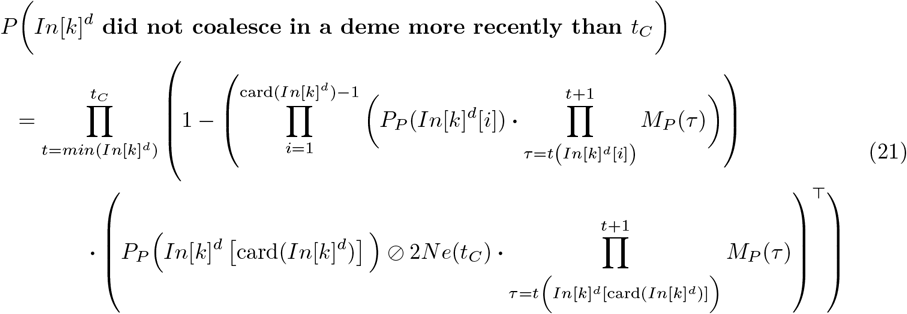

Then, the probability that the descendants *In*[*k*]^*d*^ coalesce at time *t*_*C*_ = *t* (*In*[*k*]) (or the probability of *In*[*k*]) is:

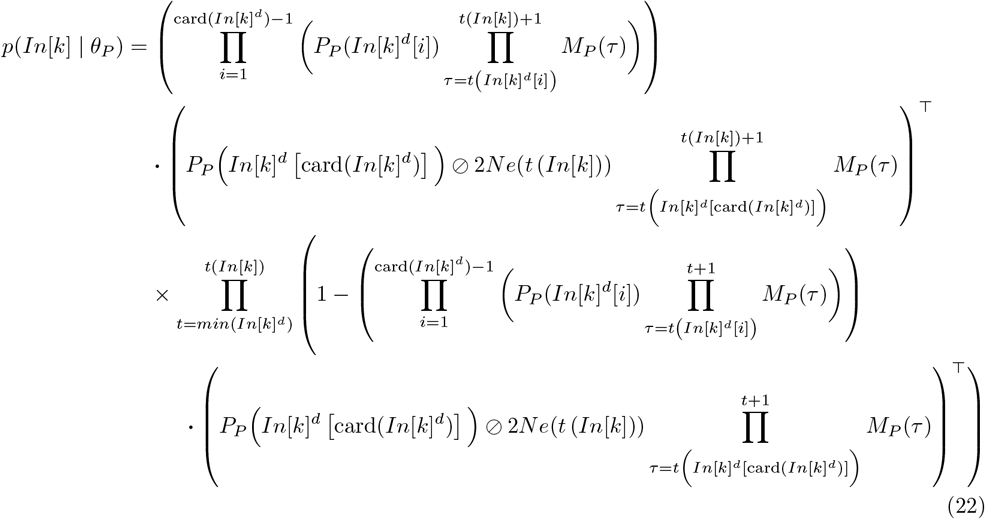

The allele probability distribution of coalescent ancestors as is :

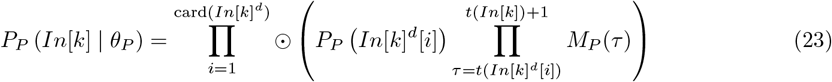

Where ⊙ is the elementwise multiplication operator

As we presented before for the probability of a genealogy knowing genetic data and model, the probability of a genealogy knowing population data and model is obtained by multiplying the probabilities of equation (22) over coalescence events and updating the probability distribution of coalescent ancestors using Equation (23) at each coalescence event:

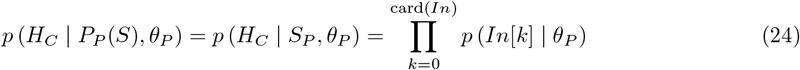

where the demographic probability distributions of the nodes *In*[*k*], used to calculate *p* (*In*[*k*] | *θ*_*P*_) using Equation (22), are successively given by Equation (23).

The algorithm to calculate the demographic probability of a genetic-simulated genealogy is as follows:

#### Algorithm 5

Calculation of the probability of a coalescent history given a genetic model

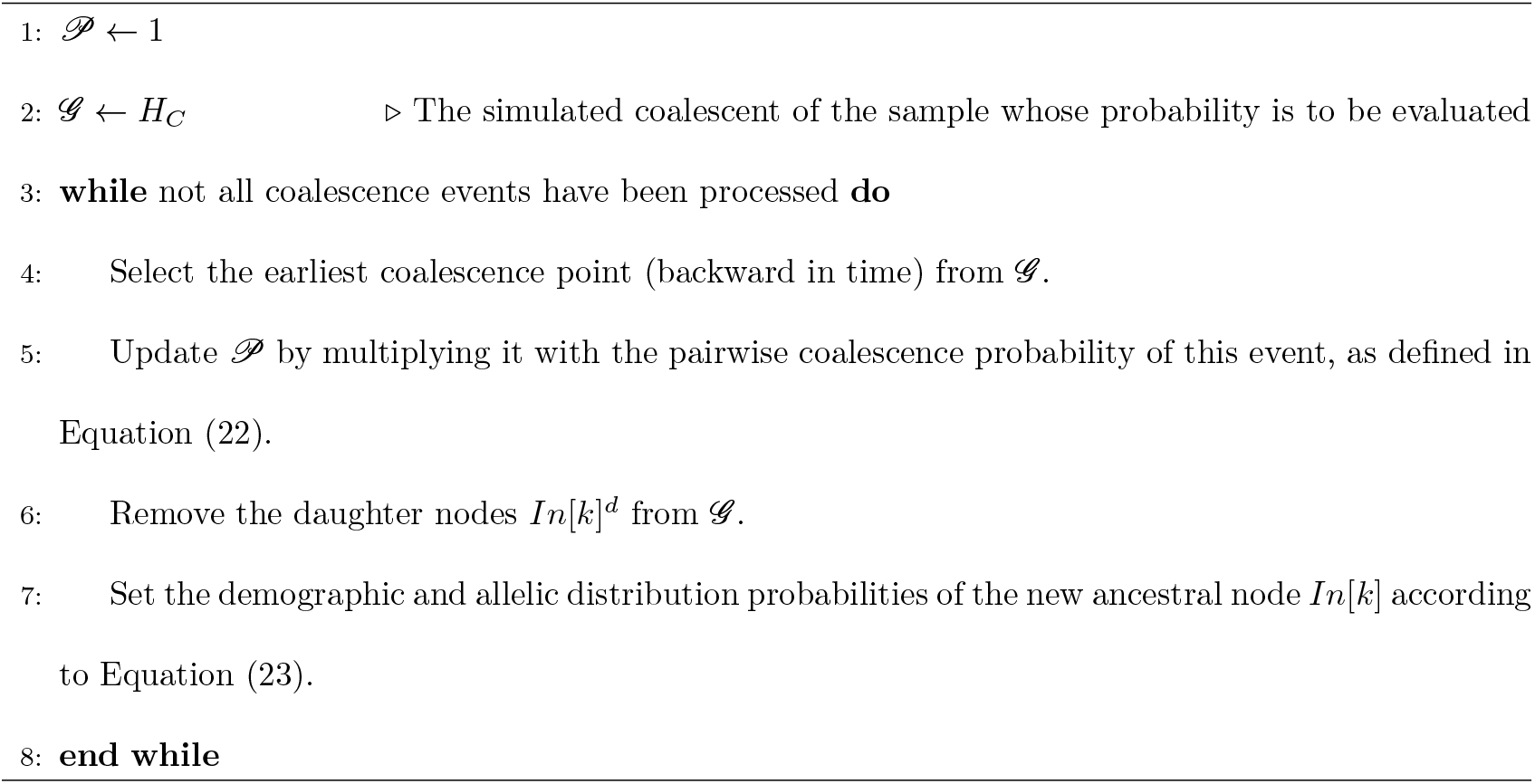

This algorithm can be combined with the genetic simulation algorithm so that the coalescence simulation directly yields the demographic probability of the genetic coalescent simulation.

#### Algorithm 6

Joint simulation of coalescence on the genetic graph and probability calculation in the population graph

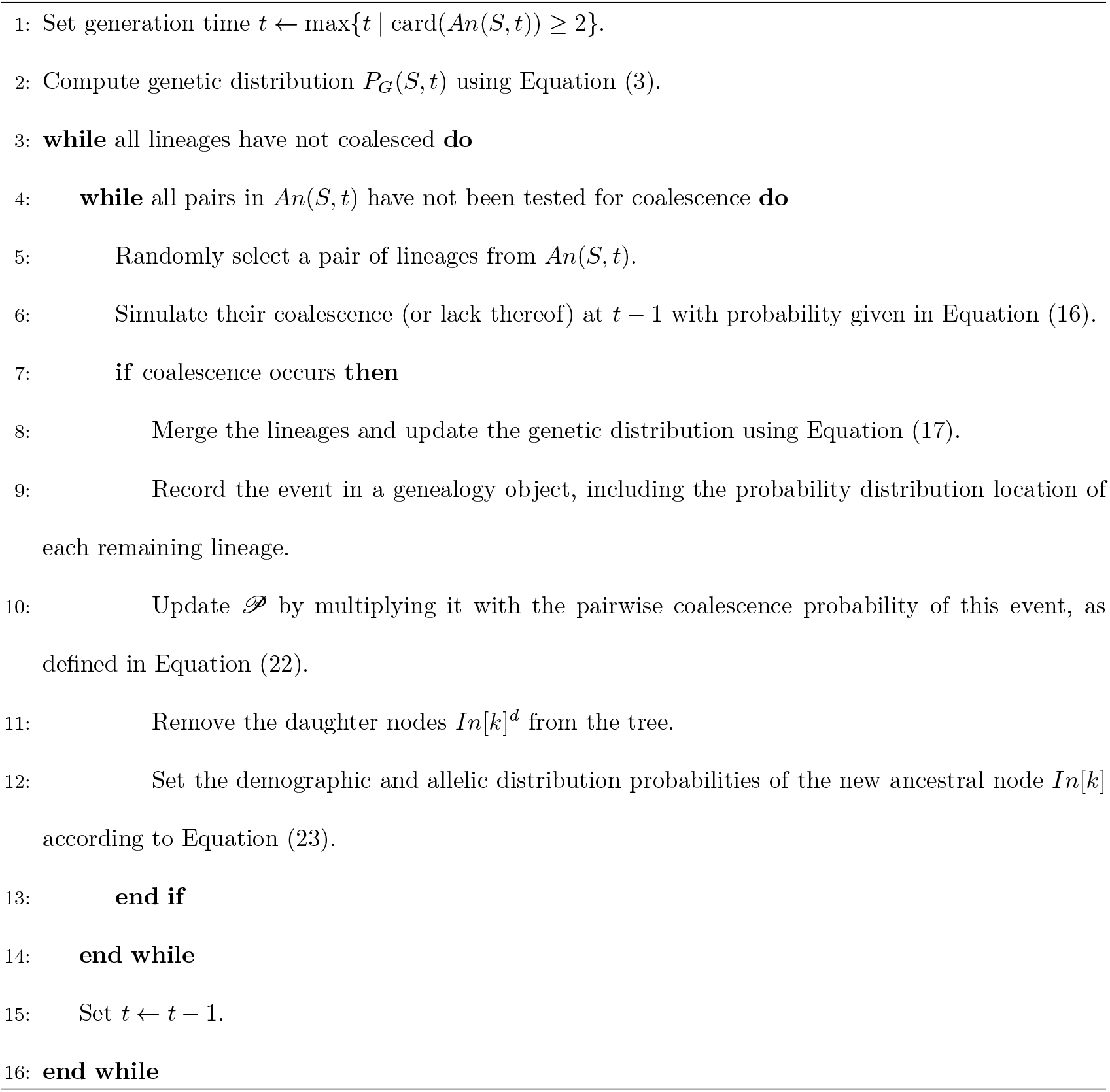

### Likelihood

The likelihood function is not analytically tractable. The coalescent is considered as latent data that can be simulated from one component of the metamodel, and its probability can be calculated using the other component. If we simulate the coalescence graph from demographic data (denoted as 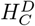) and calculate the likelihood from genetic data, we obtain:

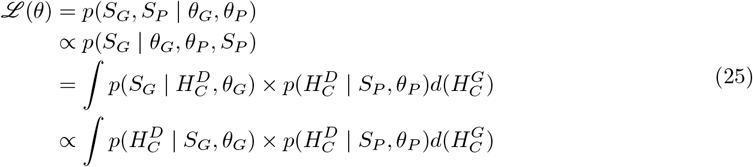

Reciprocally if we simulate coalescence graph from genetic data (we note this graph 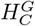) and calculate likelihood from demographic data:

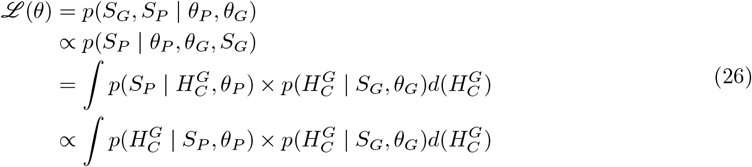

The likelihood can be estimated using a Monte Carlo algorithm, by simulating from demographic data:

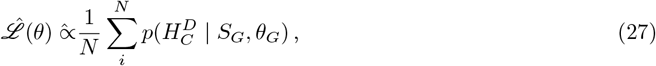

or from genetic data.

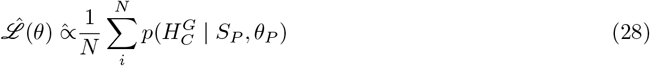

### Inference

The Monte Carlo estimated likelihood can be used in Bayesian inference or maximum likelihood estimation. For the latter, and in such a process divided into an underlying process that cannot be directly observed (in our case, the coalescent) and a sampling model (in our case, the probability of the data given the coalescent), Park [15] developed a meta-model assuming a Gaussian distribution of the log-likelihood, which allows for obtaining maximum likelihood statistics.—

### Grouping generations for computational tractability

At each generation, genes migrate across demographic and genetic space, eventually coalescing when they coexist in the same demographic group (population) and genetic group (allele). However, these events occur with low probabilities, making direct simulation computationally expensive. To reduce computation time, we can raise the transition matrix to a power, allowing for faster simulation and genealogy probability calculations.

Since information is extracted from the joint genetic and demographic distribution of the species, the effective population size *Ne*_*i*_ can be averaged within a deme over a time period corresponding to the mutation timescale (typically more than 10^4^ generations for most genetic markers). If the environment is changing, the potential effective population size *Ne*_*i*_ should be approximated using the harmonic mean of potential effective sizes over that period.

To determine an appropriate time-scaling factor, simulations should be conducted while accounting for population size, migration rates, and mutation rates.

## Discussion

In this paper, I propose a method to efficiently calculate a metamodel likelihood function for estimating genetic parameters (such as mutation rates) and environmental demographic parameters (such as the relationships between demogenetic factors—effective migration and effective population size—and social-ecological environmental variables) from population genetic data and measurable social-ecological history variables.

This work represents the first full integration of fundamental niche processes and realized niche processes related to historical movement in demogenetics. Additionally, it introduces the first likelihood-based approach in environmental demogenetics, opening pathways for maximum likelihood and exact Bayesian inference.

Demogenetic models are inherently complex, and integrating niche and historical processes adds further challenges. Likelihood-based inference has been restricted to simple coalescence models where the likelihood function is tractable. To address the computational challenges of complex “real-life” demogenetic models, Approximate Bayesian Computation (ABC) was introduced [16] and has become the primary approach for spatial demogenetic modeling [17]. Yet, ABC-based inference relies on summary statistics, whose selection is ambiguous [14, 18], and where acceptance rates decrease with an increasing number of statistics, making estimator consistency difficult to maintain [18, 19].

Instead, we employ direct likelihood-based inference. To reduce computational burden, we integrate the likelihood calculation within the simulation process—either incorporating genetic probability calculations into demographic simulations or vice versa. This approach can be further optimized by grouping demographic generations using a population transition matrix at time scales comparable to mutation time scales.

Studies must be performed to determine the identifiability of demographic parameters from genetic distributions. Since effective population size depends significantly on fluctuations, the rate of recovery may provide signals on population structure [20]. An exhaustive model may incorporate growth and carrying capacity parameters as functions of environmental data. Alternatively, to reduce computational burden we may assume a constant intrinsic growth rate (*R*) while allowing the carrying capacity (*K*) to depend on environmental data. Finally, a further simplification can be made by assuming that the species always maintains its effective population size at the carrying capacity.

From a population genetics perspective, genetic structure was historically related to simple drift, migration, mutation, and selection—forces model. With the advance of genetic sequencing and computation technologies, theses later forces were progressively linked to complex landscapes [21]. Demogenetics has emerged as a field to infer historical demography models from genetic data [17]. The integration of demogenetics with environmental niche models, termed iDDC (integrated Distribution, Demographic, and Coalescent modeling) [6], has been attempted. However, previous efforts primarily coupled independent distribution, demographic, and coalescent models rather than forming a truly integrated framework. The novel approach we propose jointly infers niche and dispersal parameters within a single comprehensive model. In this framework, the niche model predicts effective population size and migration, the key variables of interest in population genetics.

From an ecological perspective, modern studies increasingly combine biogeography, paleontology, genetics, and ecological modeling to explain species distributions [22]. Species presence is determined by a complex interplay of historical constraints (evolutionary history, dispersal, past climate) and current ecological conditions (abiotic and biotic factors). Predicting the realized niche—the fundamental goal of ecology—requires considering fundamental niche constraints, species interactions, historical dispersal, and adaptation. Our approach improves prediction by incorporating genealogical factors and dispersal constraints into distribution modeling using molecular genetic data.

Traditional species distribution models infer niche parameters from occurrence data, but this inference is biased because distribution data reflects not only niche constraints but also dispersal limitations and historical processes. By integrating niche inference with genealogical and dispersal models, we can better distinguish deterministic and stochastic processes underlying species occurrence. This integration removes biases in niche parameter estimation: in predicting effective population size, we account for coalescent-based probabilities of species presence. Although environmental niche potential is a necessary but not sufficient condition for species presence, our coalescent-based approach helps validate whether this niche was realized as predicted by migration models. This integration of dispersal and niche processes could also improve forecasting of future distributions under environmental change scenarios, including climate and land-use shifts.

A key limitation of our model is that it does not yet account for species interactions, niche evolution, or adaptation. We assume that the relationship between species density and environmental variables remains constant over time. However, since our niche models are inferred through genealogical processes, they may capture niche range across evolution in the niche model. Social-ecological data could incorporate anthropological and biotic factors beyond abiotic constraints, enabling a more comprehensive ecological modeling framework. A natural next step would be to infer multiple species’ demogenetic models simultaneously, allowing for better representation of species interactions.

We present a method to efficiently compute a likelihood function for any social-ecological demogenetic model that can be represented as a connected graph. This advancement enhances the integration of ecology and genetics, leveraging genetic and environmental data to infer population demographic scenarios. Our contributions include:

- The first likelihood-based inference method for demogenetic models in heterogeneous landscapes, previously limited to ABC approaches.
- A method that directly infers niche parameters rather than merely mapping demographic parameters, overcoming previous limitations.
- An algorithm capable of modeling diverse social and ecological network structures (spatial grids, social groups, ecological interactions), broadening the applicability of genetic inference models.

This algorithm is currently being implemented in the R library graphPOP [23]. The availability of large genetic datasets—particularly for agricultural species, pests, diseases, and endangered species—combined with hypotheses on social-ecological drivers of genetic variation, presents a significant opportunity. Our method has the potential to enhance population genetics as a tool for biodiversity and epidemiology studies in agriculture ecology and conservation by enabling more precise demographic and ecological inferences.

By fully integrating potential and realized niches within a genealogical framework, and considering that genetic data, in this respect, contain more information on ecological parameters than simple occurrence data, this approach enables a more accurate understanding of species distributions and their future projections.

## Acknowledgments

This work has been supported by IRD EGCE research unit project and BASC labex. I thank to Camille Coron and LMO Orsay France Laboratory, and Arnaud Becheler and Lacey Knowles laboratory members at of Ecology and evolutionary Biology department of Michigan Sate University for fruit full discussions.

